# LIPEA: Lipid Pathway Enrichment Analysis

**DOI:** 10.1101/274969

**Authors:** Aldo Acevedo, Claudio Durán, Sara Ciucci, Mathias Gerl, Carlo Vittorio Cannistraci

**Affiliations:** Biomedical Cybernetics Group, Biotechnology Center (BIOTEC), Center for Molecular and Cellular Bioengineering (CMCB), Center for Systems Biology Dresden (CSBD), Technische Universität Dresden, Tatzberg 47/49, 01307 Dresden, Germany; Lipotype GmbH, Tatzberg 47, 01307 Dresden, Germany; Brain Bio-Inspired Computing (BBC) Lab, IRCCS Centro Neurolesi “Bonino Pulejo”, Messina, 98124, Italy

## Abstract

**Motivation:** Analyzing associations among multiple omic variables to infer mechanisms that meaningfully link them is a crucial step in systems biology. Gene Set Enrichment Analysis (GSEA) was conceived to pursue this aim in computational genomics, unveiling significant pathways associated to certain gene signatures under investigation. Lipidomics is a rapidly growing omic field, and absolute quantification of lipid abundance by shotgun mass spectrometry is generating high-throughput datasets that depict lipid metabolism in a plethora of conditions and organisms. In addition, high-throughput lipidomics represents a new important ally to develop personalized medicine approaches, investigate the causes and predict effective biomarkers in metabolic diseases, and not only.

**Results:** Here, we present Lipid Pathway Enrichment Analysis (LIPEA), a web-tool for over-representation analysis of lipid signatures and detection of the biological pathways in which they are enriched. LIPEA is a new valid resource for biologists and physicians to mine pathways significantly associated to a set of lipids, helping them to discover whether common and collective mechanisms are hidden behind those lipids. LIPEA was extensively tested and we provide two examples where our system gave successfully results related with Major Depression Disease (MDD) and insulin resistance.

**Availability:** The tool is available as web platform at https://lipea.biotec.tu-dresden.de.

## 1 Introduction

Many bioinformatics approaches in genomics and proteomics aims to detect omic signatures (Ciucci et al., 2017), for instance the collection of genes that significantly change under a certain biological condition or that differ in case-control studies (Huang, Sherman, & Lempicki, 2009). Normally, such omic signatures need a secondary analysis in order to be understood in biological terms and linked to significant pathways (Chagoyen & Pazos, 2011). The methodology used in such cases is called *functional enrichment analysis* and, since it was originally proposed, a hundred of variations and different implementations have been developed. Nowadays, scientists in many omic fields make intensive use of enrichment analysis tools such as, to name a few, GSEA in genomics (Subramanian et al., 2005), MPEA (Kankainen, Gopalacharyulu, Holm, & Orešič, 2011) and MBRole (Chagoyen & Pazos, 2011) in metabolomics and, GeneTrail2 (Stöckel et al., 2016) in multi-omics (transcriptomics, proteomics, miRNomics, genomics).

Lipidomics is an emerging field that aims at the large scale identification and quantification of diverse lipid repertoire in biologic systems that play critical roles in cellular functions (Gross & Holcapek, 2014; Sales et al., 2016). Although lipidomics is not the most developed omic field, its importance is increasing constantly over the years, particularly nowadays that absolute quantification methods by shotgun mass spectrometry are becoming widely available (Shevchenko & Simons, 2010). Therefore, we have developed and implemented a free and open web platform called LIPEA (Lipid Pathway Enrichment Analysis) that can automatically detect the pathways and categories that are significantly associated to the multiple lipid signature provided by the user.

LIPEA web platform is available at *https://lipea.biotec.tu-dresden.de*. During the development of LIPEA, a special attention was given to design a tool that is user-friendly and offers an advanced usability. Indeed, the user interface was built on modern and auto-adaptive (responsive) web technologies, allowing the free access from any device with Internet connection. Moreover, the user does not need any programming knowledge in languages such as R, MATLAB, Python, etc., and the usage of the web tool is very intuitive and requires few trial to be easily managed.

Here, we introduce LIPEA and its interface for functional analysis of lipid signatures at the system level. LIPEA works with ID of lipid compounds contained in the Kyoto Encyclopedia of Genes and Genomes (KEGG Database; Ogata et al., 1999) and finds significantly perturbed pathways, applying statistical tests. LIPEA adopts the Fisher exact test, where the probability that the random event could happen is given by the hypergeometric distribution. It was extensively tested and below we provide two examples related with Major Depression Disease (MDD) and Insulin resistance.

## 2 Methods

The architecture of LIPEA was implemented adopting the model-view-controller (MVC) pattern that has been pursued for a clear design which separates different responsibilities within an interactive application (Veit & Herrmann, 2003). To simplify the use of this approach, we built our system using *Symfony Framework*, which is a stable and documented tool in this field (https://symfony.com/). Symfony allowed us to develop a modular platform, with a high degree of abstraction that provides a remarkable scalability, allowing the addition of new modules and components in the future. The idea behind this architecture is to identify specific altered pathways - provided by the KEGG Database - using exclusively lipid compounds. The approach used to this task is the Over Representation Analysis (ORA) (Church, Tavazoie, Hughes, Campbell, & Cho, 1999; Draghici, Khatri, Martins, Ostermeier, & Krawetz, 2003).

### 2.1 Architecture

The MVC pattern achieves independence by decoupling data access, dataprocessing logic, data presentation and user interaction tasks into three distinct object classifications (Curry & Grace, 2008). These classifications are represented in different layers: Model layer (blue), Controller layer (orange) and View layer (green) respectively (Fig. 1). Moreover, Symfony facilitates the creation of highly flexible solutions and is prevalent in systems that must provide multiple views of the same data.

**Fig. 1.**
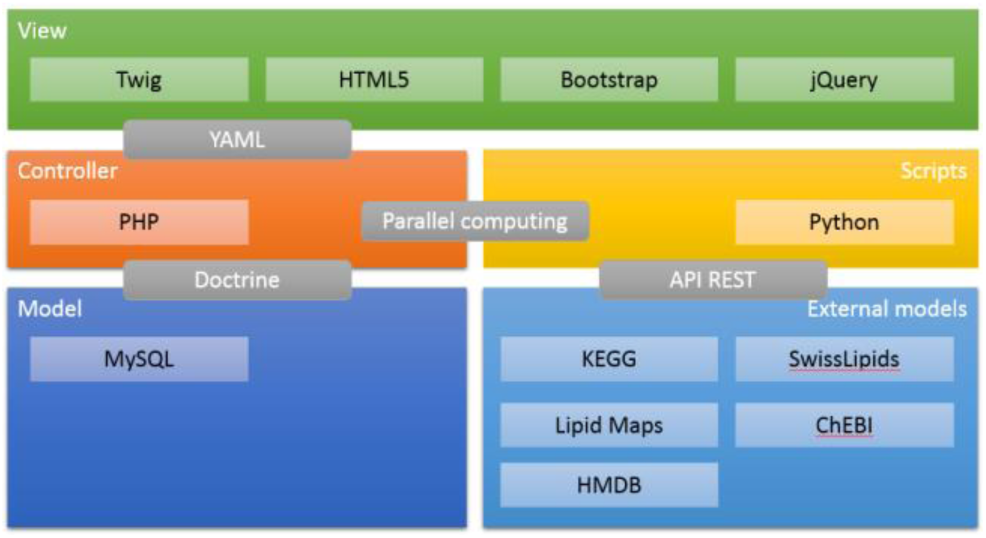
LIPEA MVC architecture. The colored boxes represent the MVC layers (Model layer [blue], Controller layer [orange] and View layer [green]), while the grey boxes repre-sent the connectors between layers.

**Fig. 2.**
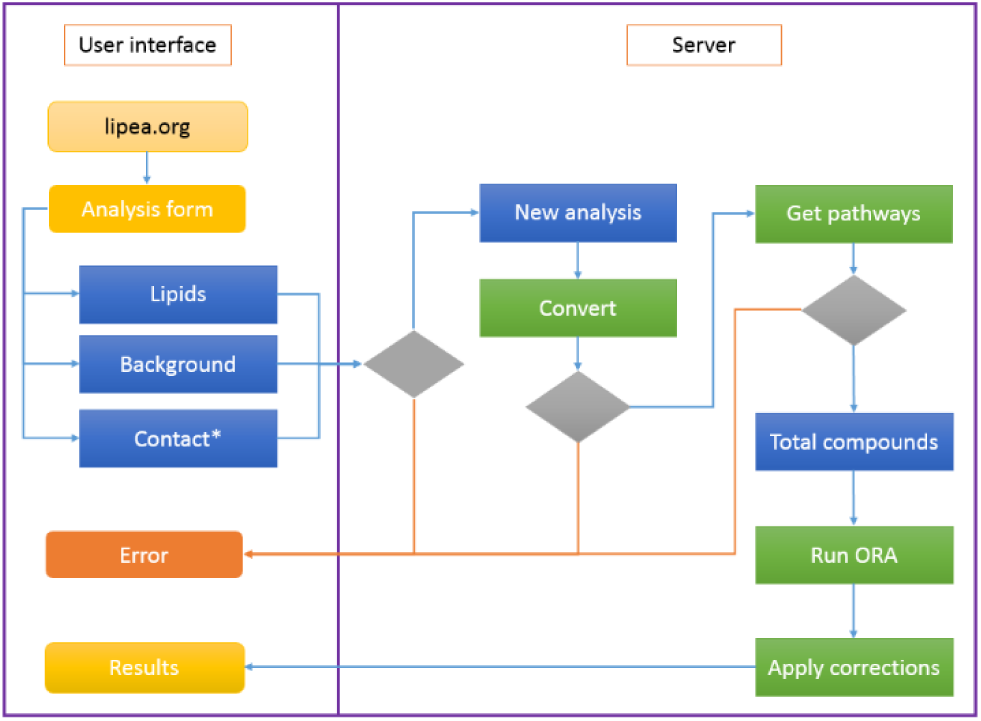
LIPEA workflow. Here two big areas are represented; 1) The user interface (left side) and 2) the server (right side). The yellow (and the orange one) boxes represent the web pages visualized by the user. The blue boxes correspond to variables, while the green ones are the procedures executed by the server. Finally, the grey rhombuses represent de-cision nodes. (*) The contact information is optional.

The View layer contains all the web pages shown by LIPEA to the user. A current problem with many bioinformatics web tools is its design and usability. Many of them are not compatibles with small screen devices (such as smartphones, tables, etc.), hindering their usability and user-interaction. To avoid this problem, we implement our interface on Twig, HTML5, Bootstrap and jQuery. Bootstrap is crucial in our design, because this tool implements an automatic screen adaptation, allowing access to our platform from any device, without losing its usability and interaction capabilities. Moreover, jQuery and its component AJAX allow asynchronous queries to our server. This gives the possibility of carrying out multiple processes where, if there are running processes, a new process can be executed without having to wait for the previous ones to finish.

The logic of this platform is contained in the Controller layer. This layer encapsulates the interaction between the views and the model. An important task of this layer is to run the analysis using parallel processes. For this approach we created separated Python scripts to run ORA and the pvalue corrections; in this case with two alternatives: Benjamini (Benjamini & Hochberg, 1995) or Bonferroni-Holm correction (Holm, 1979). So, when the user submits the analysis request and the pathways recognition is initiated, in the next steps all the calculations are execute concurrently in multiple processes, in order to reduce considerably the execution time. This layer is also connected with the externals models where we obtained a universe of IDs from multiple related databases. For this task we created a *mapping process*, that generates a table containing the relationships of all the IDs from KEGG, Swiss Lipids (Aimo et al., 2015), Lipid Maps (Fahy, Sud, Cotter, & Subramaniam, 2007), ChEBI (Hastings et al., 2013) and HMDB (Wishart et al., 2007). Given these relationships, we can extract any ID from a particular database (e.g. Swiss Lipids, Lipid Maps, ChEBI or HMDB) to obtain the corresponding KEGG ID for identifying then the associated pathways (mapping process).

Finally, the model is organized using MySQL (https://www.mysql.com/) as database system. However, we used Doctrine (http://www.doctrine-project.org/) as connector between the controller layer and the model layer. This component lets a model have multiple controllers, which can be created and altered independently from the model. Moreover, we can generate changes in the database (alter tables, create tables, add attributes, etc.) or change the database from inside the system, without the need to modify the source code of the application.

### 2.2 Workflow

The analysis starts with a form separated in four steps (Fig. 3). Step 1) *Lipids*: here the users can paste (or upload) the list of lipids that compose the signature identified in their study, and for which they want now to perform a functional enrichment analysis. Step 2) *Background*: in this step the users can define the background list and the specific organism. The background list is the original list of lipids from which the tested lipid signature was derived. Step 3) *Contact*: this step is optional, here the users can give an email address for receiving a link with the results. Step 4) *Submission*: finally the users sends the information to the server to start the analysis.

**Fig. 3.**
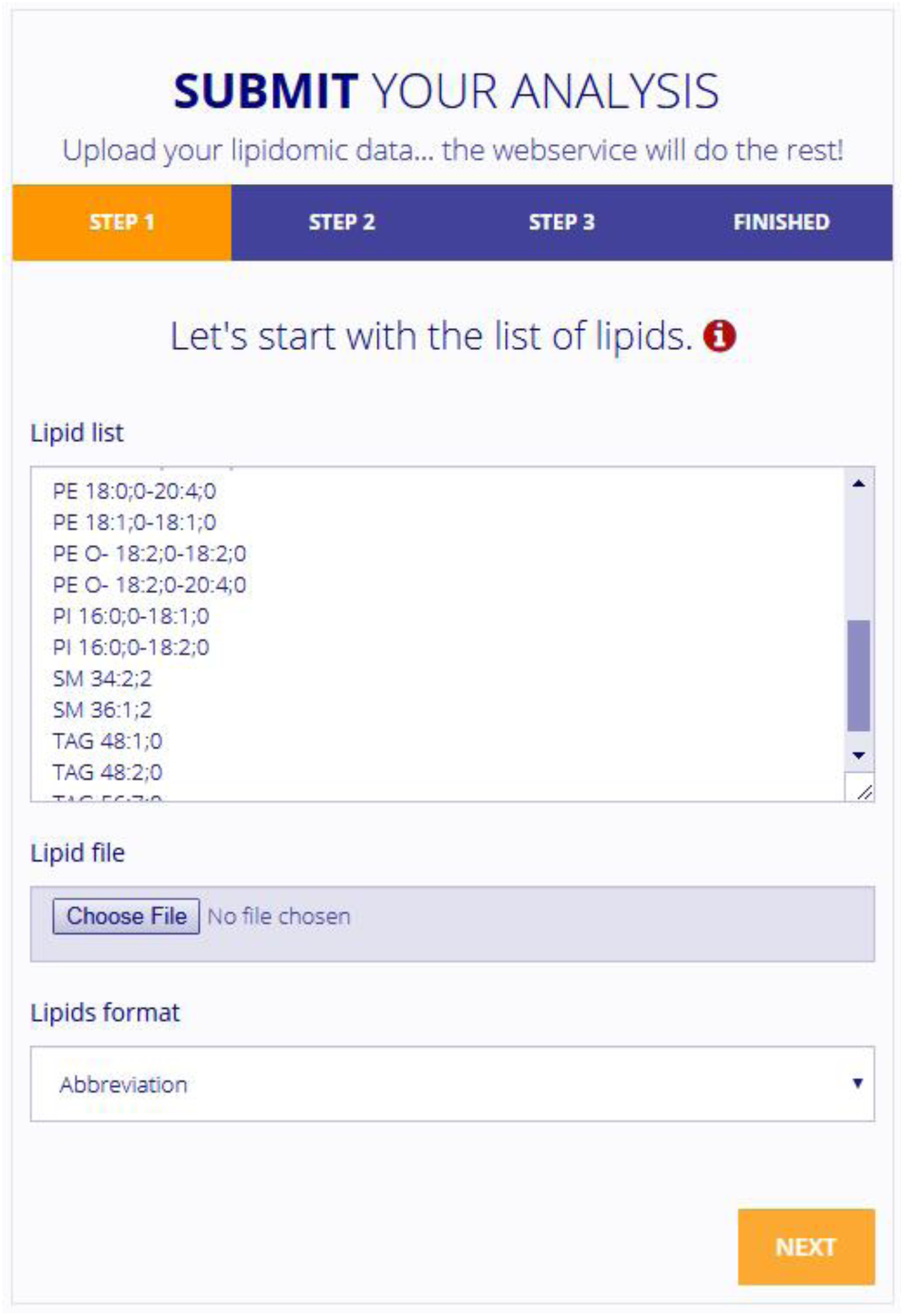
Screenshot of the LIPEA input interface. This interface is separated in four steps: Step 1) Lipids, Step 2) Background, Step 3) Contact, and Step 4) Submission, as described in the main text.

At the moment of the analysis submission, the server checks the inputs. If there are information not related with lipidomic data, the server will return an error (this verification is to avoid attacks to our server like SQL injection, among others). Instead, when the information is valid, the analysis starts and the lipid list and the background are transformed to KEGG IDs, using the internal mapping process (connected to Swiss Lipids, Lipid Maps, ChEBI, HMDB and KEGG databases via API REST). Once obtained the KEGG IDs, the server searches the pathways for the selected organism. Then, the total lipid compounds from all the pathways are extracted and the Over Representation Analysis (ORA) starts in parallel for each pathway. When all the ORA analysis are completed, the server computes the Benjamini and Bonferroni *p*-values corrections. Once this process is finished, the server returns a list of enriched pathways sorted by *p*-value. Finally, the results are shown in an interactive table, where the user can change the order, view the conversion history, among other features, and at the end download the list of pathways (Fig. 2).

### 2.3 Over Representation Analysis (ORA)

ORA starts with considering a list of annotated lipids (e.g. a lipid set related with a signature), then uses the Fisher exact test to verify if the annotations are over represented among a label (pathway) compared to the whole universe of lipids (background), which in our platform can be selected as “predefined” for a specific organism (it means, LIPEA will take all the compounds from the pathways related with the selected organism) or be a custom list given by the user (because it was the original lipid list from which the tested signature was obtained).

The steps of the algorithm used by LIPEA to implement the ORA are the following: (Step 1) Set an organism, collect the lipid list and the background. (Step 2) Select a pathway to start with. (Step 3) Tally the following 4 numbers: *m, N, k*, and *n*, where *m* is the total number of lipids in the pathway, *N* is the total number of lipids from all the pathways related with the selected organism, *k* is the number of lipids of the intersection between the lipid list and a pathway, and *n* is the total number of lipids in the list. (Step 4) Perform a Fisher exact test, with the 4 numbers obtained in the preview step, as follows.

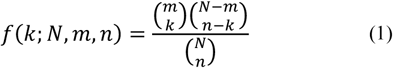

In (1) the *f* value is the probability that this random event could happen under the hypergeometric distribution. In this case, to obtain the *p*-value associated to each pathway detected by the algorithm, the following formula is used.

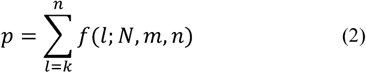

(Step 5) Go to step 2 for another pathway of interest until all are tested (Kanehisa, 2013). (Step 6) Correct the *p-*values with Benjamini or Bonferroni-Holm corrections.

## 3 Results

We tested our system with a list of potential lipid markers of major depressive disorder (MDD) (Liu et al., 2016). The authors proposed a combinational lipid panel including LPE 20:4, PC 34:1, PI 40:4, SM 39:1,2, and TG 44:2 as potential diagnostic biomarkers for MDD. As inputs we used these potential biomarker as lipid list, together with a default background for *Homo sapiens (Human)*, this means that all the lipids included in all the pathways for this organism were considered as initial lipid list from which the signature was derived.

We obtained one significant pathway as results: *glycerophospholipid metabolism* (KEGG code: hsa00564) with Benjamini and Bonferroni corrected *p*-value less than 0.05 (Fig. 4). This results have coincided with the discovery and validation of plasma biomarkers for MDD proposed by the same authors (Liu et al., 2015) where they revealed that some biomarker lipid classes related with the *glycerophospholipid metabolism* are increased in MDD subjects. However, while LIPEA automatically detected the association with the *glycerophospholipid metabolism* and proved that is statistically significant, in order to arrive to the same conclusion Liu et al. had to perform an extensive expert-based evaluation of the literature and their conclusion was not supported by any statistical test that quantifies the significance of the association between the lipid marker classes and the pathway. This example clarifies how LIPEA offers an automatic tool that emulates the ability of an expert to detect meaningful associations between lipid signatures and molecular mechanism, but with the advantage also to provide statistical significance to the proposed functional annotation, which is produced in few seconds or minutes (in relation with the query load).

**Fig. 4.**
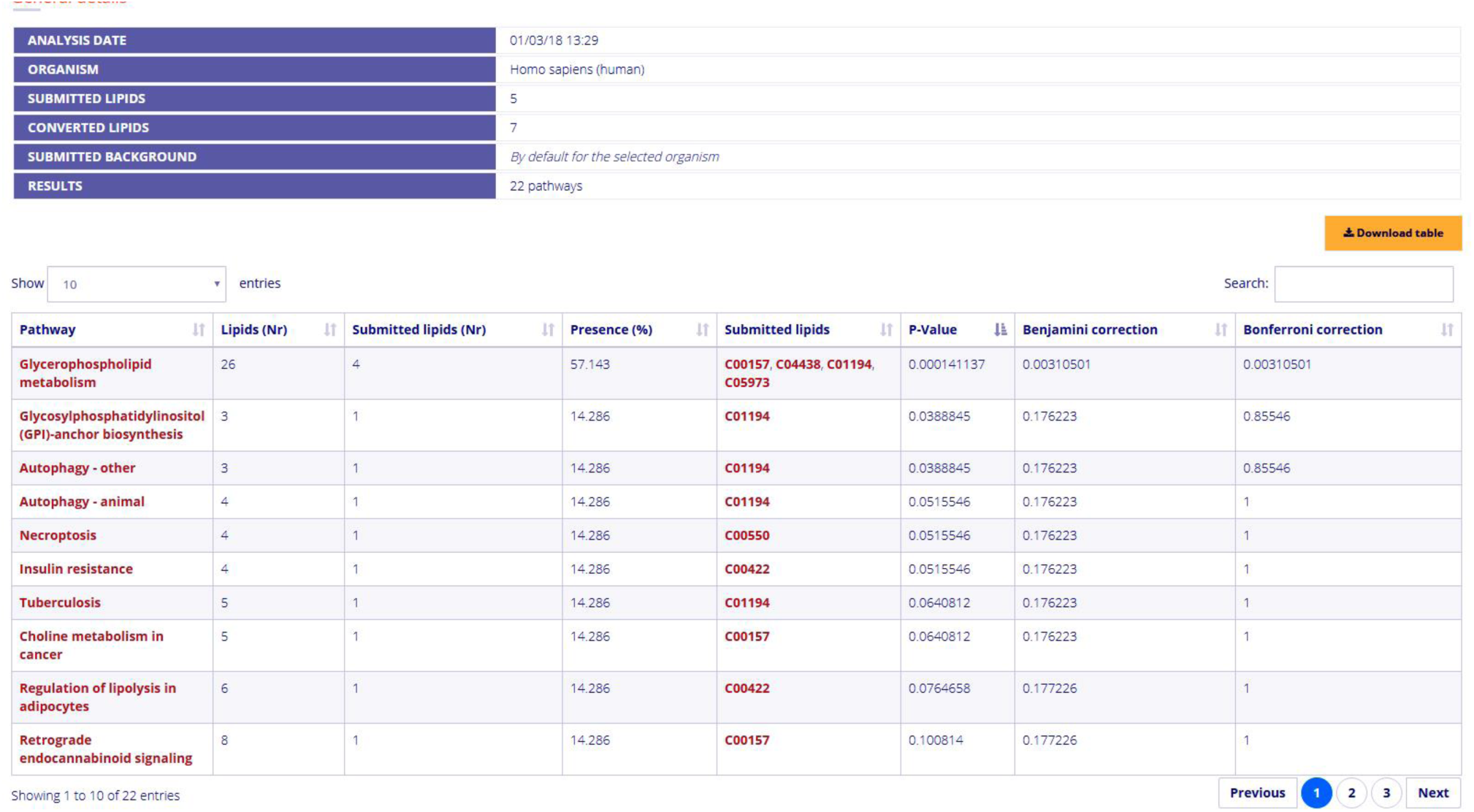
LIPEA results interface. List of results from the lipid markers LPE 20:4, PC 34:1, PI 40:4, SM 39:1, 2, and TG 44:2 related with Major Depression Disease (MDD). Our system gave as significant pathway glycerophospholipid metabolism that according with the literature is increased in MDD subjects.

Finally, to provide a second example, we took as inputs a specific list (*n* = 23) of lipids all derived from the same lipid class (*Triacylglycerides*, TAG) and *Rattus norvegicus (Rat)* as default organism for the background. LIPEA identified as significant pathway *insulin resistance* (KEGG code: rno04931), and in agreement with this result we found studies where the authors proposed that an enhancing of triglycerides is related with the insulin sensibility in rats (Lee et al., 2006; Pinnamaneni, Southgate, Febbraio, & Watt, 2006; Schenk & Horowitz, 2007; Todd, Watt, Le, Hevener, & Turcotte, 2007). This second example clarifies how LIPEA could be used as tool not only for test but also for hypothesis generation. In fact, if - in absence of experiments - we make a theoretical hypothesis on the relevance of a set of lipids for a certain molecular mechanism, by means of LIPEA we can mine the support that our hypothesis can have according to the knowledge currently available in the databases.

## 4 Discussion

Given as inputs three terms (the lipid signature of interest; the original background list of lipids; and the organism under investigation) LIPEA runs automatically in seconds the entire functional enrichment analysis, providing in output a list of pathways in which a sub-set of the tested lipids are over-represented. If the background list is not uploaded by the user, the list of all the available lipids (in the LIPEA’s database) for the selected organism is used instead. Finally, the updated list of pathways used in LIPEA for a selected organism can be downloaded by the user in case of necessity. LIPEA currently support all the organisms available in KEGG database. Moreover, our platform has an administration system where all the information about the organisms, pathways and compounds are updated continuously.

Currently LIPEA analyses can only be done at the class scale. Although there are some databases like Swiss Lipids, Lipid Maps, etc. that have information at the (sub)-species level, this information is not directly related with the pathways contained in KEGG. Moreover, the lipidomic field at the moment does not offer an adequate knowledge for pathways that is at the lipid species level, because the technology that analyzes samples at the species level in a robust and replicable manner is becoming largely available to the scientific community only recently (Han, Yang, & Gross, 2012; Shevchenko & Simons, 2010; Subramaniam et al., 2011; Yetukuri, Ekroos, Vidal-Puig, & Orešič, 2008). However, we would like to clarify that this is not a limitation of LIPEA itself but of the current data available on the public databases.

Given the above, in the future we aim to develop the LIPEA project in order to integrate the new knowledge generated in lipidomics at the scale of species and sub-species. This will aim to perform a transversal enrichment analysis identifying the significant pathways at multiscale level (classes, species, and sub-species).

## Acknowledgements

We thank *Lipotype GmbH* (https://www.lipotype.com/) for providing us the lipid classes used to build our mapping and some images for the web platform. The authors also gratefully acknowledgment Alexandre Mestiashvili, systems administrator in Prof. Michael Schroeder group in Biotechnology Center (BIOTEC) Technische Universität Dresden (TUD) for his support during the system deployment.

## Funding

This work was supported by: the independent group leader starting grant of the Technische Universität Dresden (TUD) and the Klaus Tschira Stiftung (KTS) gGmbH, Germany (Grant number: 00.285.2016). Claudio Duran was supported by the DAAD Forschungsstipendien Promotionen fellowship.

## Contributions

C.V.C. envisaged the study. A.A. and C.D. implemented the source code. A.A. developed the web platform with suggestions of S.C., C.D., C.V.C., and M.G. A.A. and

C.D. created the IDs lipid mapping table with the main help of M.G. A.A. created the figures. A.A., C.D. and C.V.C wrote the article with inputs and corrections of S.C., and M.G. C.V.C. led, directed and supervised the study. All authors tested the web platform and have read and approved the manuscript.

### Conflict of Interest

none declared.

